# Adverse perinatal outcomes for obese women are influenced by the presence of comorbid diabetes and hypertensive disorders

**DOI:** 10.1101/559856

**Authors:** Evelyne M. Aubry, Stephan Oelhafen, Niklaus Fankhauser, Luigi Raio, Eva L. Cignacco

## Abstract

Maternal obesity often occurs together with comorbid diabetes and hypertensive disorders. All three conditions are independently associated with negative perinatal outcomes. Our objective was to determine the risk and burden of adverse perinatal outcome that could attributed to maternal obesity in combination with a comorbid status.

We analyzed data from 349’755 singleton deliveries in Switzerland between 2005 and 2016. For the association of maternal obesity in the presence or absence of comorbidities with various perinatal outcomes, we estimated adjusted relative risk (RR) using multivariate regression modeling and determined the multivariate-adjusted attributable fraction (AFp).

Regardless of comorbidities, obesity was a main predictor for macrosomia, fracture of the clavicle, plexus paresis, failure to progress in labor and prolonged labor. However, we identified a second subset of outcomes, including neonatal hyperglycemia and preterm birth, that was only significantly linked to obesity in the presence of comorbidities. A third subset of outcomes was independently influenced by either obesity or comorbidities.

We suggest that comorbidities such as diabetes and hypertensive disorders should be considered when relating maternal obesity to adverse perinatal outcomes.

## Introduction

Obesity is one of the greatest health problems globally and is considered a major cause of death and disease in industrial countries^1^. The rise of epidemic obesity is also reflected in an increased prevalence of maternal obesity. In the US, more than 50% of pregnant women are considered obese, while in Europe, the number of women considered overweight and obese during pregnancy has also increased significantly, attaining rates of 30-37% in 2010^2,3^. One third of these women were considered obese^4^. According to a study by Frischknecht at *al.*^5^, the prevalence of women with obese pre-pregnancy BMI in Switzerland almost doubled between 1986 and 2004.

Obesity, defined as a BMI ≥ 30 kg/m^2^, has been shown to impact mothers’ and infants’ health by increasing the risk of adverse perinatal outcomes such as stillbirth^6^, congenital malformations^7^ and delivery complications^8^. The rise in obesity prevalence is considered to be the major determinant of the striking increase in obesity attendant comorbidities in pregnancy, mainly gestational diabetes and hypertensive disorders^9^. The term comorbidity refers to a situation in which one or more disorders co-occur in the same individual, either at the same time or in some causal sequence^10^. Many women are diabetic, hypertensive or both, in addition to being obese. This indicates the existence of a metabolic syndrome. In pregnancy, obesity in combination with comorbidities like pre-existing diabetes, gestational diabetes mellitus (GDM), pre-gestational hypertension, gestational hypertension as well as pre-eclampsia can present with a mixed symptom picture, modulated by these diverse comorbid conditions and thus complicate a precise diagnosis. Furthermore, the interaction between illnesses can worsen the course of both, the obese and the comorbid condition or adversely influence the course of the other condition.

Obesity, diabetes and hypertension have all been discussed independently regarding poor perinatal outcomes^11–13^. Each of these conditions can contribute to adverse perinatal outcomes and might worsen as part of the normal physiological changes in pregnancy. As with maternal obesity, pregnant women with pre-existing diabetes or GDM are at increased risk of pregnancy loss, perinatal mortality, fetal macrosomia and congenital malformations^14^. A review of 55 general US population studies demonstrated that hypertensive disorders during pregnancy are also a risk factor for caesarian delivery (41.4%), preterm birth (PTB) (28.1%), neonatal unit admission (20.5%), and perinatal death (4%)^15^. The complex relationship and the interplay between obesity and its attendant comorbidities and the relative contributions of each to the increased risk of obstetric perinatal complications are still unclear. Despite strong claims for the significant association of BMI and various adverse perinatal outcomes, some inconstancy between studies remains^16–18^. The recent umbrella review of 156 meta-analyses provided strong evidence for an association between obesity and only three obstetric outcomes: cesarean delivery (RR 2.00, 95% CI 1.87-2.15), preeclampsia (RR 4.14, 95% CI 3.16-4.75) and low Apgar score (RR 1.29, 95% CI 1.23-1.36)^19^, while other outcomes like fetal death, macrosomia, preterm birth were not clearly linked to obesity. Such inconsistencies may be attributable to the effects of comorbidities on the relative risk for adverse perinatal outcomes, in addition to obesity as a risk factor alone.

The objective of the current study was to investigate the interplay between obesity and its attendant comorbidities, diabetes and hypertensive disorders and how these conditions contribute to poor perinatal outcomes. We estimated the adjusted relative risk (RR) for obese women based on the presence or absence of comorbidities and thus quantified an attributable fraction of the population (AFp)^20^. While RR offers an estimate of the strength of an association, AFp considers both the RR and the prevalence of the exposure in the population. It provides information on the population disease load that is due to the underlying exposure of maternal obesity and diabetes and hypertensive disorders. It is essential to estimate the proportion of disease burden that could be prevented by reducing maternal obesity and its attendant comorbidities ^21^.

## Methods

### Study design and data collection

The present study retrospectively analyzed anonymized data from women in Switzerland who delivered singleton infants between 22 and 43 weeks gestation from January 1, 2005 to December 31, 2016.

The study utilized a computerized database containing details of deliveries collected prospectively by a Swiss obstetric study group (Arbeitsgemeinschaft Schweizerischer Frauenkliniken, Amlikon, Switzerland) during a 12-years period (January 2005 – December 2016)^22^. The group collates and manages data from more than 100 obstetrics hospitals of various sizes and structures. The quality of the data recorded was ensured by a two-steps control system. Firstly, the completeness and exactness of all data was verified at each participating center at the time of discharge by a senior obstetrician. Secondly, the plausibility of all data entered in the database was assessed by the data center quality control group. In case of data discrepancy, the hospitals were asked to verify and eventually correct the information previously given. Moreover, the risk of data entry error was reduced to a minimum by the fact that all variables included in the database, with the exception of maternal age and weight, birth weight, and umbilical cord pH, were collected as categorical variables (e.g., second stage of labor longer than two hours, maternal hemorrhage greater than 1000 mL). Items in the database contain the International Statistical Classification of Diseases and Related Health Problems, 10^th^ Revision (ICD-10) codes. In compliance with Swiss Federal Law on data protection (Human Research Act Article 2), data were anonymized and irreversibly de-identified to protect patient, physician and hospital privacy. Because data was retrospective, pre-existing, and de-identified, this study did not need approval from the ethics committee. All procedures performed were in accordance with the ethical standards of the institutional and/or national research committee and with the 1964 Helsinki declaration and its later amendments or comparable ethical standards.

### Exposures

Pre-pregnancy weight status was based on the body mass index (BMI), calculated as weight (kg) divided by height (m) squared and registered at the first prenatal visit by the physicians in charge. The physicians further categorized pre-pregnancy BMI by obesity status, defined as non-obese (BMI < 30 kg/m^2^) or obese (BMI ≥ 30 kg/m^2^) according to the World Health Organization’s definition^23^.

Maternal comorbidities according to hospitalization diagnoses were extracted from the database using the following (ICD-10) codes (in brackets): pre-existing diabetes mellitus I/II (treated) (E14.9) and gestational diabetes mellitus NOS (O24.4), whereby mothers were only counted in one of the diabetes categories. Hypertensive disorders included gestational hypertension without significant proteinuria (≥ 140/90) (O13), pre-eclampsia (O14, O14.1), eclampsia (O15.9), pre-existing hypertension (O10.9, O11).

### Perinatal outcomes

Labor outcomes included instrumental vaginal delivery methods (vacuum extraction and forceps) (O81), cesarean delivery (primary, secondary and elective section) (O82), induction of labor (physical, systemic/vaginal prostaglandin use), prolonged labor (prolonged first stage >12h, prolonged second stage >2h) (O63.0, O63.1), failure to progress in labor > 2h (O63.9), obstructed labor due to shoulder dystocia (O66.0), fetal heart rate anomaly (O68.0), epidural anesthesia, perineal injury (grade I-IV, uterine rupture, laceration of cervix and vagina wall, other trauma) (O70.0, O70.1, O70.2, O70.3, O71.1, O71.3, O71.4, O71.9), abnormal third stage of labor (retained placenta and membranes without hemorrhage, postpartum hemorrhage, obstetric shock, postpartum clotting disorder (O72.1, O72.3, O73.0, O73.1, O73.4, O75.1). Neonatal outcomes included macrosomia defined as a birth weight of 4000 g or above in babies delivered after 37 weeks of gestation. A 5 minutes Apgar scores ≤ 7 and umbilical cord pH < 7.1 were defined as abnormal^24,25^. Other neonatal outcomes were transitory neonatal hypoglycemia (P70.4), respiratory distress of newborn (P22.9), plexus paresis (p14.9), fracture of the clavicle (P13.4). Preterm birth was defined as delivery < 37 weeks of gestation. Intensive care unit admission was assumed if the infant was subsequently transferred to the pediatric unit after birth. Intrauterine fetal death (O36.4), stillbirth and infant death up to seven days post-partum (P95) were defined as fetal death.

### Statistical analysis

The prevalences of obesity, diabetes and hypertensive disorders in pregnant women across the time span of 2005 – 2016 were calculated. All analysis was performed using R version 3.4.1. Binomial logistic regression models provided adjusted relative risks (RR) and 95% confidence intervals for the association of maternal obesity in combination with comorbidities and various complications during pregnancy and delivery. Multivariate Poisson regression models were used for outcomes with a prevalence ≥10%^26^. All models were adjusted for maternal age, parity, history of smoking during pregnancy, and ethnicity. Instrumental vaginal delivery, induction of labor, prolonged labor, perineal injury, shoulder dystocia, fracture of the clavicle were additionally adjusted for caesarian section. Macrosomia was additionally adjusted for preterm birth. Outcome variables were compared to those of non-obese women or non-obese women without comorbidities as the reference group. Regression models were implemented using generalized linear models (GLM) from the stats package available in *R*^27^. P-values less than 0.05 were considered significant.

The population attributable fraction (AFp) was estimated for the effect of obesity and/or comorbidities on all obstetric outcomes using the prevalence of disease in the total population and the prevalence of disease in those with the risk factor. For each outcome the following standard formula was applied^20^:

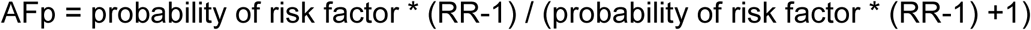

Nonparametric bootstrap confidence intervals from the *boot* package available in R was employed to model AFp while incorporating appropriate levels of uncertainty to account for variance in the estimates^28^. A thousand replicate samples were created from the original sample using with-replacement sampling. The replicate samples are used to create a 95% confidence interval (CI) defined as the 2.5^th^ and 97.5^th^ percentile of the distribution of the 1000 possible values^29^.

### Data availability

The datasets analyzed during the current study are available from the corresponding author on reasonable request.

## Results

A total of 349,975 singleton births over 22 weeks of gestation were recorded in the ASF Database between 2005 and 2016. Complete data, both sociodemographic and clinical, including in particular labor and neonatal outcomes, were available for 349,755 women (99.9%). The clinical characteristics of the study population are depicted in Table 1. A total of 25,374 (7.3%) women were classified as obese This sample of 63% Swiss, 28% European and 9% non-European matched the origin statistics of the representative sample of adult female in Switzerland (Table 1)^30^.

**Table 1.** Maternal characteristics, perinatal (labor and neonatal) outcomes (n=349755). Data are mean ± SD or count (%)

|  |  |
| --- | --- |
| **Maternal characteristics** |  |
| Age (years) | 31.02 ± 5.09 |
| Parity |  |
| 1 | 168,508 (48.2) |
| 2 | 125,392 (35.9) |
| 3+ | 55,833 (15.9) |
| Origin |  |
| Switzerland | 220,735 (63.1) |
| European | 98,524 (28.2) |
| Non-European | 30,360 (8.7) |
| Smoking during pregnancy | 21,226 (6.07) |
| Pre-pregnancy BMI ≥ 30kg/m<sup>2</sup> | 25,374 (7.5) |
| - with comorbidities | 5,246 (1.5) |
| Pre-pregnancy BMI < 30kg/m<sup>2</sup> | 324,381 (92.7) |
| - with comorbidities | 19,936 (5.7) |
| **Labor outcomes** |  |
| Instrumental vaginal delivery | 40,318 (11.5) |
| Induction of labor | 6,7467 (19.3) |
| Cesarean section | 100,483 (28.7) |
| Epidural anesthesia | 95,724 (27.4) |
| Fetal heart rate abnormality | 80,826 (23.1) |
| Prolonged labor | 33,950 (9.7) |
| Failure to progress in labor | 22,307 (6.4) |
| Abnormal third stage of labor | 25,017 (7.1) |
| Shoulder dystocia | 2,353 (0.7) |
| Perineal injury | 148,497(42.5) |
| **Neonatal outcomes** |  |
| Macrosomia | 29,550 (8.5) |
| Small for gestational age | 14,831 (4.2) |
| 5' Apgar score ≤ 7 | 18,811 (5.4) |
| Umbilical cord pH<7.1 | 13,211 (4.0) |
| Neonatal hypoglycemia | 2,627 (0.8) |
| Respiratory distress of newborn | 14,535 (4.2) |
| Fracture of the clavicle | 643 (0.2) |
| Plexus paresis | 257 (0.1) |
| Intensive care unit admission | 5,030 (4.3) |
| Fetal death | 1,474 (0.4) |
| Preterm birth <37 weeks of gestation | 19,819 (5.7) |

Obese women had a higher probability of having at least one comorbidity compared to non-obese women (21.1% in obese vs. 6.5%) (Table 2). Their pregnancy was four times as likely to be complicated by hypertensive disorders (8.4% vs. 2.2%). Obese women were treated three times more often for GDM (13.5% vs. 4.2%) than their non-obese peers. A similar proportion of obese women without a comorbidity and non-obese women without comorbidities was observed, while the prevalence for obese women that had at least one additional comorbidity was reduced by a factor of four (Table 2).

**Table 2.** Rates of comorbidities (number and %)

|  | Total<br>(n=349,755) | Non-obese<br>(n=324,381) | Obese<br>(n=25,374) |
| --- | --- | --- | --- |
| =1 comorbidity | 240,040 (6.9) | 19,389 (6.0) | 4,651 (18.3) |
| >1 comorbidities | 2,272 (0.7) | 1,560 (0.5) | 712 (2.8) |
| Diabetes mellitus type1 and 2 | 2,528 (0.7) | 1,956 (0.6) | 572 (2.3) |
| Gestational diabetes mellitus (GDM) | 17,006 (4.8) | 13,572 (4.2) | 3,434 (13.5) |
| Hypertensive disorders | 9,173 (2.6) | 7,035 (2.2) | 2,138 (8.4) |

In Table 3 adjusted relative risk (RR) and population attributable fraction (AFp) of adverse perinatal outcomes are presented for obesity in the total population. As expected, obesity was significantly associated with all adverse pregnancy outcomes that were assessed in this study except for prolonged labor, fetal death and preterm birth. The RRs were highest for hypertensive disorders (RR 3.97), diabetes mellitus type 1 and 2 (RR 3.83), and GDM (RR 3.25). Moreover, obesity contributed most to the burden of the following outcomes: hypertensive disorders (15.3%), diabetes mellitus type 1 and 2 (15.0%), GDM (13.3%), plexus paresis (8.7%), macrosomia (6.3%), and fracture of the clavicle (4.4%) (Table 3).

**Table 3.**
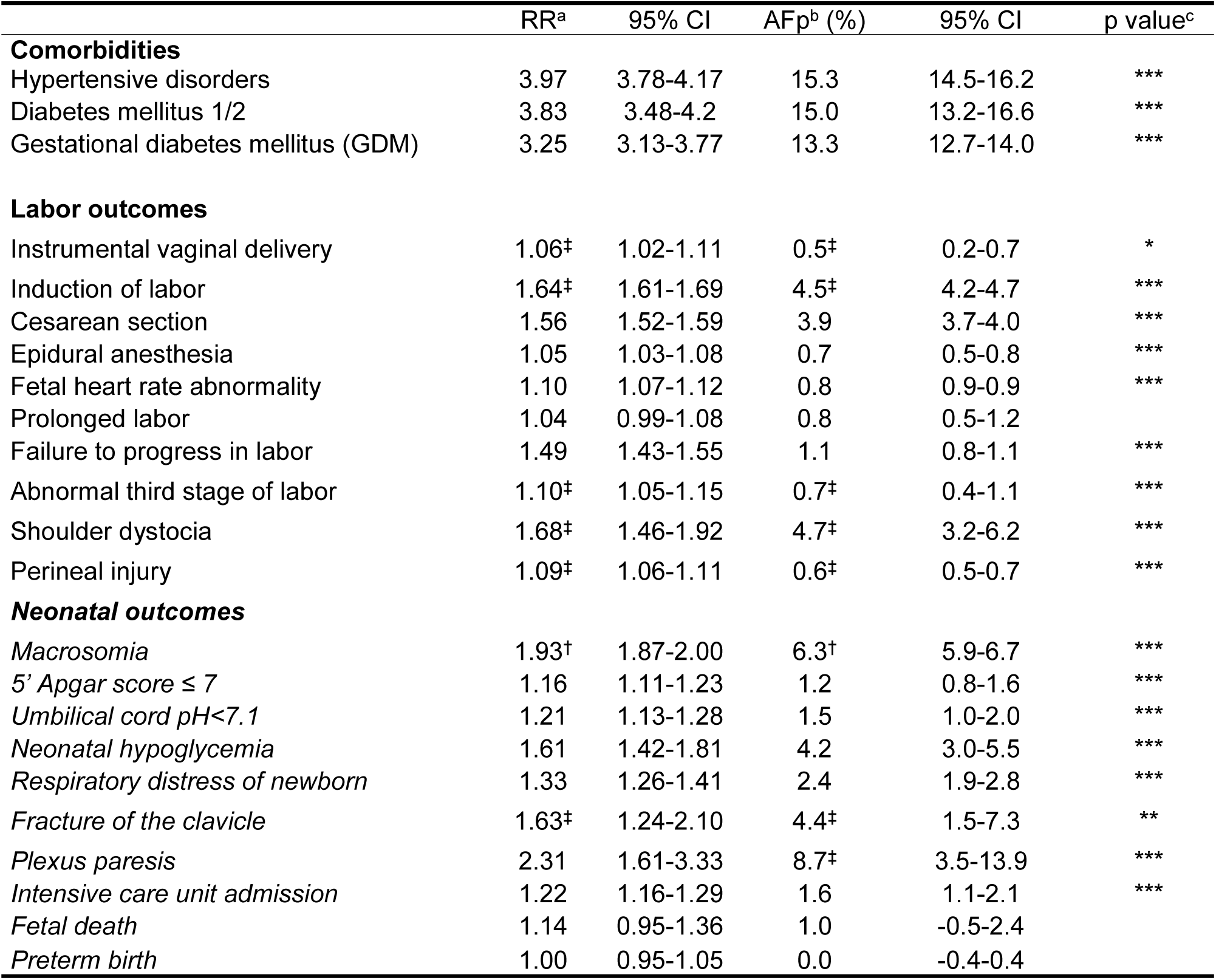
Multivariate analysis of adjusted relative risks for perinatal (labor and *neonatal*) outcomes, and attributable fraction in the population for risk factors in obese compared to non-obese women. ^a^ Poisson (prevalence >10%) or binominal (prevalence <10%) models adjusted for age, ethnicity, parity and history of smoking during pregnancy. ^b^ Attributable fraction in the population adjusted for age, ethnicity, parity and history of smoking during pregnancy. ^‡^ additionally adjusted for cesarean delivery. ^†^ additionally adjusted for preterm birth. ^c^ *pvalue<0.01, **pvalue<0.001, ***pvalue<0.0001.

### Perinatal outcomes associated with obesity

RRs and AFps were stratified based on the presence or absence of obesity attendant comorbidities and obesity (Table 4). Our results show that RRs of macrosomia in obese women almost double regardless of comorbidities diagnosed. A larger fraction of macrosomia cases could be attributed to obesity alone than to obesity in presence of comorbidities or to morbidities of non-obese mothers (Table 4). A similar pattern of association for women only affected by obesity was observed for plexus paresis, fracture of the clavicle, failure to progress in labor, prolonged labor, instrumental vaginal delivery and epidural anesthesia. Furthermore, the RRs were slightly lower for these outcomes when obese women were additionally affected by a comorbidity.

**Table 4.**
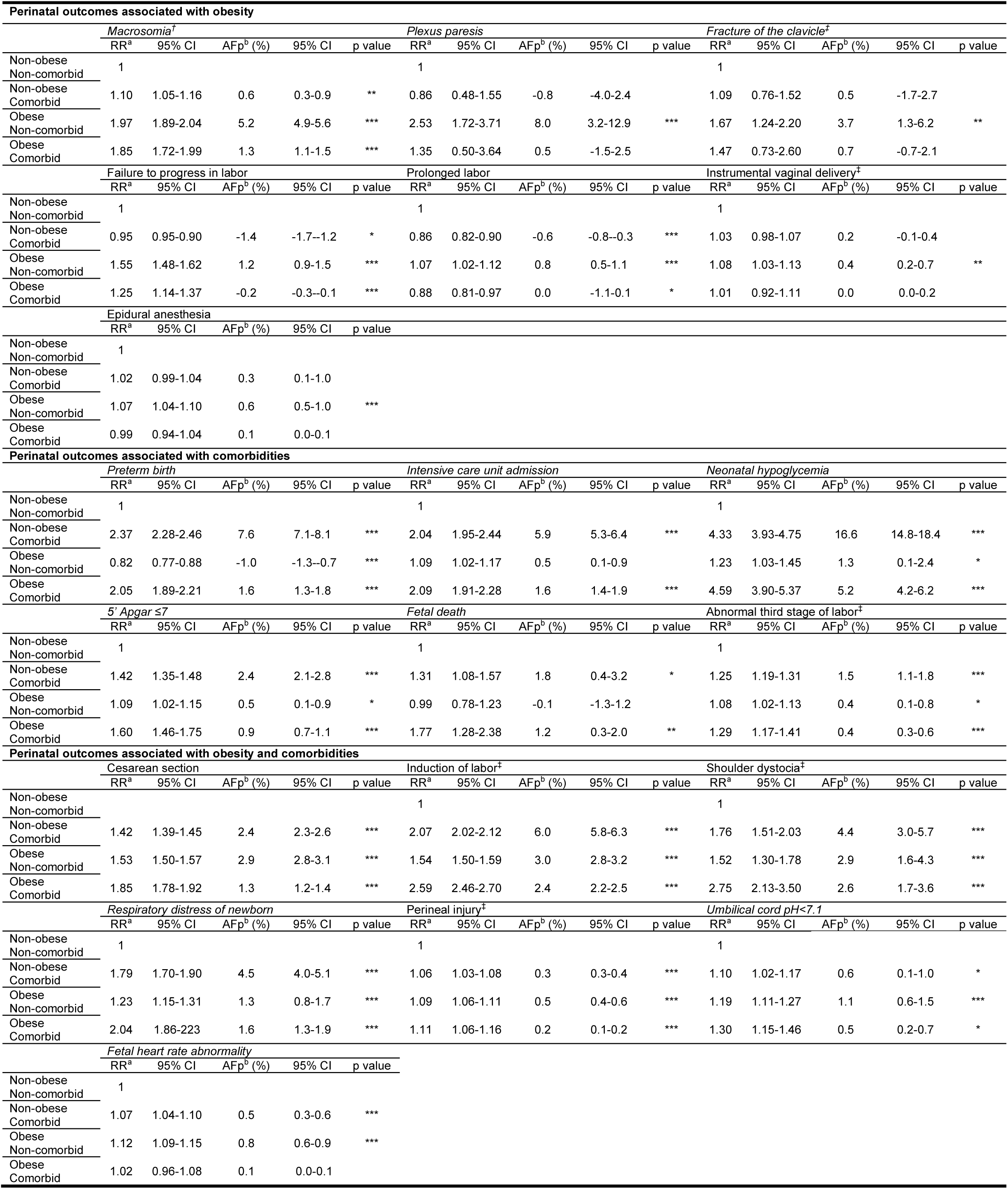
Multivariate analysis of adjusted relative risk for perinatal (labor and neonatal) outcomes, and attributable fraction in the population for risk factors with a variable combining obesity and comorbidities. A Poisson (prevalence >10%) or binominal (prevalence <10%) models adjusted for age, ethnicity, parity and history of smoking during pregnancy. ^b^ Attributable fraction in the population adjusted for age, ethnicity, parity and history of smoking during pregnancy. ^‡^ additionally adjusted for cesarean delivery. ^†^ additionally adjusted for preterm birth. ^c^ *pvalue<0.01, **pvalue<0.001, ***pvalue<0.0001

### Perinatal outcomes associated with comorbidities

In contrast, RRs and AFps for another subset of outcomes including preterm birth, intensive care unit admission, neonatal hypoglycemia, 5’ Apgar score <7, fetal death, and abnormal third stage of labor were highest when obese women suffer from comorbidities. Little or no risk and a minimal disease burden could be attributed to exclusively obese women. Non-obese women with comorbidities showed a similar significant pattern of risk and contribution to adverse perinatal outcomes than their obese peers. This suggests an association with a comorbid, rather than with an obese status.

### Perinatal outcomes associated with obesity and comorbidities

A last subset of outcomes such as cesarean delivery, induction of labor, shoulder dystocia and respiratory distress of newborn among others was independently associated with both obesity and comorbidities. Highest RRs were identified for obese women suffering from comorbidities.

## Discussion

Using population-based data of women and their offspring from Switzerland, we examined the impact of obesity and its attendant comorbidities including diabetes and hypertensive disorders on perinatal outcomes.

Our results confirm that a link exists between obesity and the risk of adverse perinatal outcomes and match with the findings of a recent meta-analysis who identified maternal BMI over 30 as a strong or at least highly suggestively associated risk factor for fetal macrosomia, low Apgar score, instrumental vaginal delivery, gestational diabetes mellitus and pre-eclampsia ^19^.

Maternal obesity, diabetes and hypertensive disorders seem to be closely linked and often occur in the same patient. They also independently increase the health risk for mother and fetus/infant^31^. Despite the indications of interaction, little is known about the relative risk contribution of maternal obesity, and comorbid diabetes and hypertensive disorders to adverse perinatal outcomes. We showed that macrosomia, plexus paresis, fracture of the clavicle, failure to progress in labor and prolonged labor are significantly associated with maternal obesity, but were only slightly influenced by comorbidities. Similarly, a study found maternal hyperglycemia and obesity to be independent predictors for various adverse perinatal outcomes^32^. By demonstrating distinct higher risks for overweight women, with or without GDM, they confirm the relevance of increased weight independently of hyperglycemia. This might be explained by the fact that diabetes and hypertensive disorders are manageable components during pregnancy.

Not only does glycemic control in diabetes or antihypertensive treatment reduce the impact of these comorbidities, but they may even be beneficial for the biological mechanisms of parturition in obese women^33^. Indeed, we observed that risks for macrosomia, failure to progress in labor and prolonged labor are slightly but significantly lower in obese women suffering from a co-morbidity, compared to obesity alone. The observed protective benefit might be due to an unidentified effect of the comorbidity treatment or, to a subtle change in compliance with a diet that accompanied a specific therapy. Obese women with GDM who achieved desired levels of glycemic control using insulin therapy had similar macrosomia rates to normal-weight controls. However, this “positive” effect was eliminated by exclusively diet-controlled therapy^34^.

In contrast, for a subset of other perinatal adverse outcomes we identified increased risks only when obese women presented additionally comorbidities. For example, we showed that obese women with comorbidities exhibit a more than two-fold risk for preterm birth. However, preterm birth was prevented when obese women did not suffer from any comorbidity. Tsur et *al.*^35^ showed a comparable protective effect of obesity on the risk of preterm birth. Stratification of obese women with and without comorbidities resulted in a decreased risk of preterm birth associated with obesity, independent of comorbidities. The authors argued that fat tissue confers protection for preterm birth through alternation of metabolic factors such as tumor necrosis factor alpha or obesity-associated gene FTO variants^35^. However, a Spanish study linking overweight to GDM, confirmed the positive correlation of glucose intolerance and pre-term deliveries, independent of the mother’s BMI, but it did not show a preventive role of obesity^32^. A possible explanation is the adjustment of preterm birth prevalence for macrosomia, which is known to be a factor linked to post-maturity^36^. Likewise, in our study, diabetes and hypertensive disorders rather than obesity appear to be exclusive risk factors for intensive care unit admission and fetal death. Even though recent reviews identified an elevated risk for intensive care unit admission (OR 1.5) and fetal death (RR 1.46) associated with obesity, they agreed that maternal obesity itself, increases the risk for comorbidities that are risk factors for stillbirth, preterm birth and subsequently intensive care unit admission^37,38^.

In the present study both obesity and comorbidities can be considered as independent risk factors for a third subset of outcomes including cesarean section, induction of labor, shoulder dystocia and respiratory distress of newborn. It has been shown in multiple studies that the excessive risk for caesarean section among women is irrespective of whether they are either obese, diabetic or hypertensive or a combination of these conditions^8,39,40^. Unrelated events can require caesarean section. On the one hand, the cephalo-pelvic disproportion of macrosomic babies of obese mothers can lead to non-progressive labor, resulting in emergency cesarean section^41^. On the other hand, increased blood pressure in the mother and preeclampsia, often resulting in intrauterine growth restriction, increases the cesarean section rates to over 50%^42,43^. The risks for cesarean section is highest for obese women with comorbidities since they encounter events due to their obesity, then again due to their comorbid status.

Finally, we took advantage of our large, comprehensive dataset to determine how the burden of obesity in combination with its comorbidities could be attributed to adverse perinatal outcomes. Generally, maternal obesity was found to be the cause for 4.2% of neonatal hypoglycemia (table 3). However, obesity alone contributed only to 1.3% of neonatal hypoglycemia, while when combined with comorbidities it increased to 5.2% (Table2). The contribution to neonatal hypoglycemia of comorbidities in obesity seems striking when we consider that in this study population, the prevalence of obese women without comorbidities was almost four times greater when compared to their peers who suffered from comorbidities (5.7% and 1.5%). This highlights why it is all the more important to consider the comorbidities of diabetes and hypertensive disorders when looking at the health burden of maternal obesity.

Nevertheless, reducing maternal obesity will not only be beneficial for preventing a subset of adverse outcomes, but positively impact all perinatal outcomes by lowering the prevalence of comorbidities. Our data show that a hypothetical elimination of maternal obesity would lead to an approximate 15% reduction in each comorbidity. With increased obesity prevalence there is even greater potential in decreasing comorbidities though obesity prevention. In Canada, where obesity prevalence is 25%, half of the GDM and almost a third of hypertensive disorders in pregnancy could be avoided by eliminating maternal obesity^44^. Consequently, up to 50% of averse perinatal outcomes associated with comorbidities would be prevented, if maternal obesity were eliminated there.

Our study took advantage of a unique standardized dataset from a large, consistently collected, representative sample of women provided by Swiss hospitals. However, the dataset was limited by the lack of additional socio-economic confounders. Nevertheless, substantial evidence to support the observed links exists and we made every effort to address major confounding factors such as age, parity, ethnicity and history of smoking in our analysis. In addition, stratifying women with comorbidities might create a selection bias^35^. Obese women suffer from diabetes or hypertension related to obesity, whereas the pathogenesis of these morbidities in non-obese patients is different, and therefore confounds the comparability between these groups. Irrespective, it does not affect the comparison between obese women and obese diabetic or hypertensive women.

## Conclusions

We showed that maternal obesity is associated with comorbid diabetes and hypertensive disorders. These further impact on a variety of adverse obstetric complications not necessarily directly linked to obese conditions, and are therefore difficult to capture in studies by investigating obesity as exclusive underlying exposure for adverse outcomes.

Obesity attendant comorbidities can act independently and should be considered when associating obesity with dysfunctional labor. From a public health perspective, we advise that interventional studies related to obesity in pregnancy should control for the effect of comorbidities. Conversely, decreasing pre-gravid obesity should have at least the same priority as any efforts to prevent and control diabetes and hypertensive disorders.

## Acknowledgements

None.

## Authors Contributions

E.A. contributed to the design of the study, the analysis of the data, the interpretation of the data and the writing of the manuscript. S.O. and N.F. contributed to the analysis of the data and critically revised the manuscript. L.R. contributed to the design of the study, to the acquisition of the data and critically revised the manuscript. E.C. contributed to the interpretation of the data and critically revised the manuscript.

## Additional information

Competing interests: The authors declare no competing interests.

